# Development of duplex real time PCR for quick detection of *Cryptosporidia* in goats

**DOI:** 10.1101/2022.02.23.481731

**Authors:** Atul Kumar Sharma, K. Gururaj, Rama Sharma, Anjana Goel, Souvik Paul, Dinesh Kumar Sharma

## Abstract

Cryptosporidium spp. is the most important foodborne and waterborne pathogens and the leading cause of mortality from foodborne and waterborne gastrointestinal disease. In neonates of domestic animals it is associated with consistent diarrhoea and dehydration. Cryptosporidium infection begins with the ingestion of sporulated oocytes disseminated by carrier animals that consistently contaminate the environment. Many diagnostic tests are available including microscopy, antigen trap ELISA, but none of the diagnostic tests available currently cannot differentiate between active and passive infection in the host. In the current study, to address this challenge an mRNA based duplex TaqMan^®^ probe PCR (dRT-qPCR) was developed to target the Cryptosporidium oocyst wall protein (COWP) gene and 18ssu rRNA gene in a single tube that can detect metabolically active Cryptosporidial oocysts. The mRNA transcripts are the direct indicator of any actively replicating cell and it will help decipher the active stages of its lifecycle in host. This diagnostic assay was standardized by computing transcript copy number-based limit of detection. For COWP and 188ssu rRNA genes the limit of detection was 7.08×10^04^ and 5.95×10^05^ respectively. During active infections the oocyst wall protein will be active and so its COWP gene transcripts will act as marker for active infection. While transcripts for 18SSU rRNA are constitutively expressing in Cryptosporidial life cycle. This current diagnostic assay will be a quantitative marker that will help assess active stages of Cryptosporidium infection in neonates. The disease dynamics will help better understand to formulate the control strategies and contain infection among the healthy animals.

**Importance:** Cryptosporidiosis is an important neonatal disease affecting goats causing diarrhoea, dehydration and stunted growth. For diagnosing this condition, many diagnostic tests are available including microscopy, immunological tests, but none of the diagnostic tests available currently can differentiate between active and passive infection in the host. The mRNA transcripts are the direct indicator of any actively replicating cell and especially in intracellular parasites it will help decipher the infective stages of a lifecycle in the host, and hence the test was developed in a reverse transcriptional format in a duplex mode. The currently developed diagnostic assay for cryptosporidiosis was evaluated for sensitivity using Limit of detection (LOD). This diagnostic test will act as a quantitative marker to aid in detecting active stages of *Cryptosporidium* infection in neonatal goats and will eventually lead to better control strategies for managing cryptosporidial infections in the future.

---

Cryptosporidiosis disease caused by the intracellular, extra-cytoplasmic protozoan parasite, belong to the phylum Apicomplexa. Cryptosporidiosis is the primary cause of chronic diarrhea among immunocompromised patients especially in children suffering from HIV, AIDS-defining illness because of its high association with mortality (1–4). *Cryptosporidium* oocysts are transmitted via faecal-oral route, either direct or indirect contact with contaminated or ingestion of contaminated water and food (5, 6). In domestic animals, the disease is economically important as it affects the growth rate due to persistent clinical diarrhoea especially in neonates. The disease mainly affects the goat kids, calves and lambs, while the adult animals act as silent carrier disseminating the infection (7, 8). The animals upon infection with *Cryptosporidium* oocysts, excystation occur and then subsequently release four sporozoites. Sporozoites adhere to the epithelial cells of ileum and incorporated into the parasitophorous vacuole, and the feeder organelle present in all intracellular stages of *Cryptosporidium* acts as interface between host-parasite cells. After feeder organelle development, sporozoite forms a trophozoite and becomes more spherical in shape, asexual reproduction begins and develops the type I meront and release merozoites, which immediately re-infect the host. Again type I meront begins the asexual reproduction and forms the type II meront, which release the four merozoites and start the sexual reproductive cycle. Merozoites capture the host cells and transform it into the macrogamonts or microgamonts, which release the microgametes and macrogametes. Microgametes fertilise the macrogametes and produces a zygote, which differentiate into four sporozoites, develops as oocysts and released in lumen and again re-infect the host from thin-walled oocysts. Thin-walled oocysts are shed in stools and act as source of infection for other hosts, and are also responsible for the auto infection in the host. Whereas, thick-walled oocysts excreted by infected host are very resistant to pH, humidity, temperature etc. and can survive several months (9–12).

Diagnosis still plays a very important role because in neonatal animals many other diseases like colibacillosis, rotaviral diarrhoea caused by group A rotavirus (GARV) and bovine corona virus (BCoV) occurs commonly and there is a need to differentiate them or identify them collectively in case of mixed infection. While coccidiosis occurs in weaned animals in later stages that might perplex in the proper diagnosis. Microscopy is an excellent tool and can be highly specific but requires expertise and experience in identification of the weak acid fast round oocysts from other artefacts similar to them. Besides conventional microscopy and staining, molecular diagnostic approaches like PCR are also gaining momentum Molecular diagnostic assays are usually regarded as more sensitive compared to microscopy and serological diagnosis in human and animals for the detection of *Cryptosporidium* in several studies (13–23). The molecular assays are determined by the factors like stage of life cycle/infection as well as method of nucleic acid extraction for targeting the *Cryptosporidium* spp. (24–27).

The already available diagnostic assays can only detect the presence or absence of *Cryptosporidium* in a qualitative manner, and there is no scope to decipher the metabolically/transcriptionally active oocysts from passive ones. Hence, a robust technique is required that can able to differentiate the active oocysts from dead oocyst in passive infection.

In detail, the main aim of this study is to develop and standardize an mRNA based *Cryptosporidium* spp.-specific real-time PCR assay using two important target genes viz. 18 small subunit ribosomal RNA (*18SSU rRNA*) and *Cryptosporidium* oocyst wall (*COWP*). The *18ssu rRNA* gene is highly abundant in terms of genomic copies and it has high transcripts that can increase the sensitivity of the assay, while the *COWP* is available only in transcriptionally active live cryptosporidial oocysts. (13-23, 28-31).

Hence, in the current study we report an mRNA based TaqMan® probe duplex real time PCR that can able to detect the metabolically active *Cryptosporidium* oocysts from dead oocysts in passively infected animals

## RESULTS

### Microscopic identification of *Cryptosporidium*

*Cryptosporidium* suspected faecal smear that were stained by the modified Ziehl-Neelsen staining (**mZN staining**) were examined by microscopy at 100x magnification under immersion oil, where 39 samples were found positive out of 61 suspected samples which are collected form neonatal goat kids. The oocysts appeared as acid fast round bodies against a blue-green background (Fig.1).

**FIG 1.**
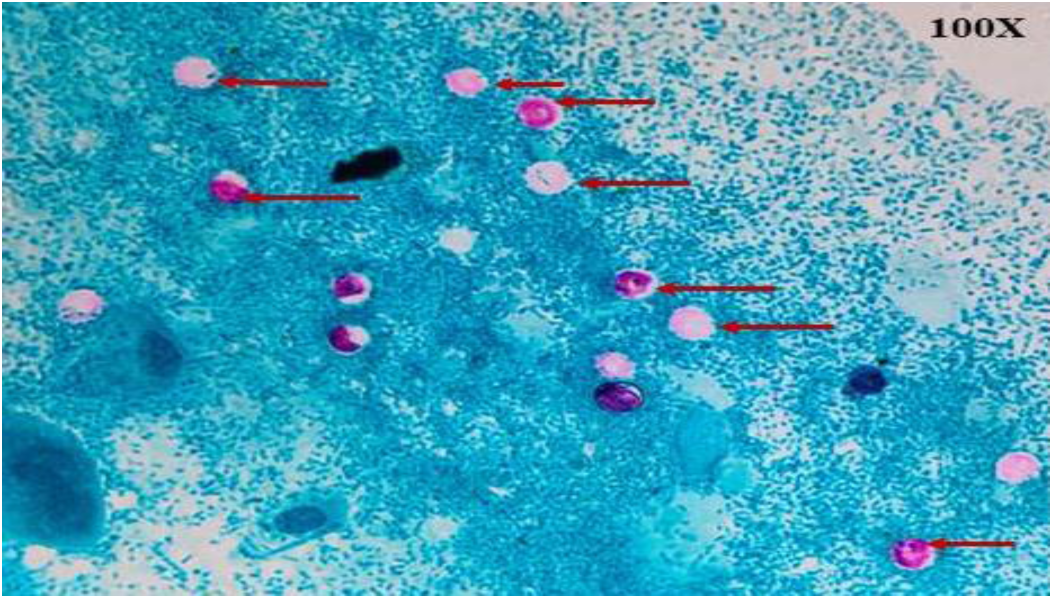
Faecal smear stained from clinically affected Cryptosporidiosis disease animal using modified Ziehl Neelsons’ technique showing *Cryptosporidium* oocysts (Indicating Arrows)

### Conventional Polymerase Chain Reaction for *COWP* and *18ssu rRNA* gene

The conventional PCR reactions for newly designed primers viz., *COWP* and *18ssu rRNA* were analysed by gel electrophoresis. The amplicon size for COWP and 18ssu rRNA is 157bp and 109bp respectively and the same was observed in the gel photo (Fig.2). Whereas the negative controls did not generate any visible amplicons in this technique.

**FIG 2.**
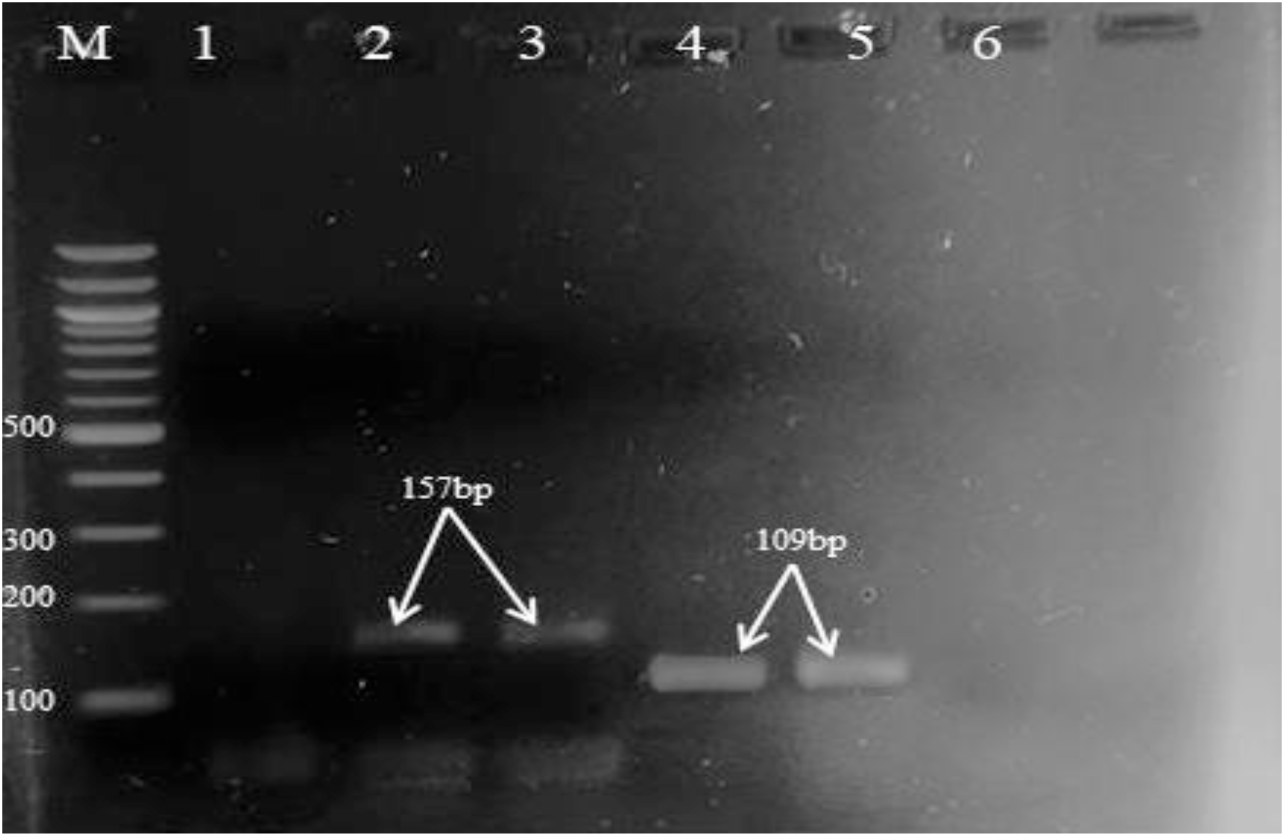
Gel-electrophoresis of conventional PCR targeting COWP and 18ssurRNA genes using newly designed primers. Lane M: 100bp ladder, Lane 1 NTC, 2&3 Positive samples *COWP -* FAM (157bp), 4&5 positive sample for*18ssu rRNA*-HEX (109bp), Lane 6: NTC (no template control).

### DNA duplex PCR for *Cryptosporidium*

After verifying the workability of the newly designed primers by conventional PCR, the same set of primers, but this time along with their respective probes was assayed using a duplex format using template DNA positive for cryptosporidium. Signals were produced in the form of Cq for both the *COWP* (FAM) and *18ssu rRNA* (HEX) as marked as visible sigmoid curve highlighted as blue and green respectively (FIG 3). The Cq and RFU values for *COWP* and *18ssu rRNA* generated in duplex DNA PCR are provided in table below (Table 7). While the no template controls (NTC) produced mean Cq for *COWP* and *18ss rRNA* of 33.32 and 35.88 respectively which were above the threshold cut-off cq for their respective genes and hence are negative.

**FIG 3.**
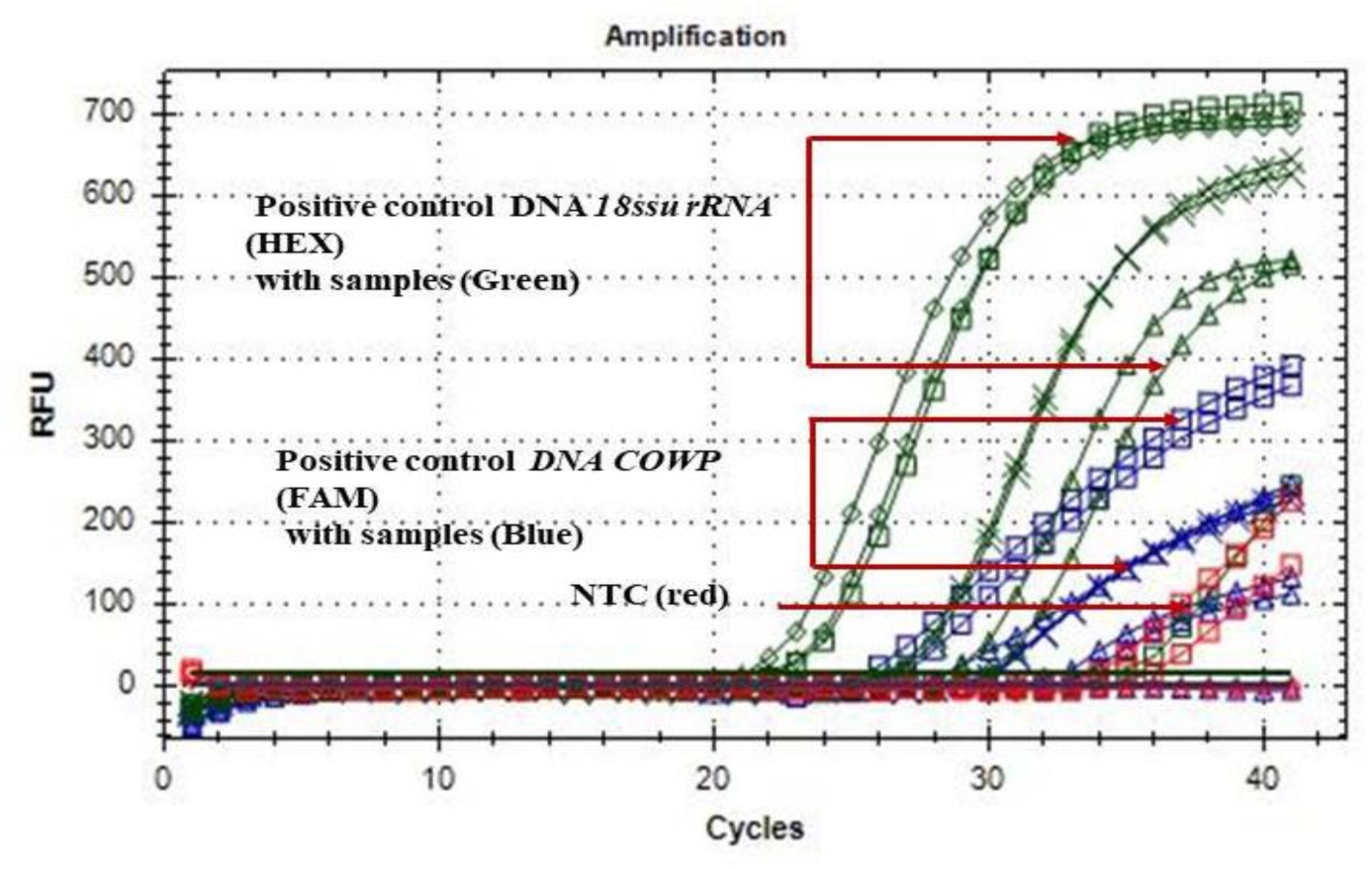
DNA duplex PCR for *COWP* (FAM) and *18ssu rRNA* (HEX) gene of *Cryptosporidium* from positive DNA with NTC.

### Probe and Primer Standardization for reverse transcription duplex TaqMan probe real time PCR

After observing the workability of the duplex TaqMan® probe real time PCR using DNA template, we further worked on the optimal concentrations of probe and primers by titration for cDNA (prepared from faecal RNA positive for *Cryptosporidium* both by microscopy and conventional PCR). Based on the titrations, the primer or probe concentration that has produced the best RFU value and the Cq values were taken as the optimal concentration. So as per the results (Fig.4), for *COWP* gene the optimal primer and probe concentrations were 8picomoles (for both forward and reverse primers) and 10picomoles respectively. Similarly, the optimal primer and probe concentrations for *18ssurRNA* are 4picomoles (each of the primers) and 4pmol for probe. Henceforth these standardized concentrations were used for assay of unknown samples suspected for *Cryptosporidium* throughout the study for the current diagnostic technique.

**FIG 4:**
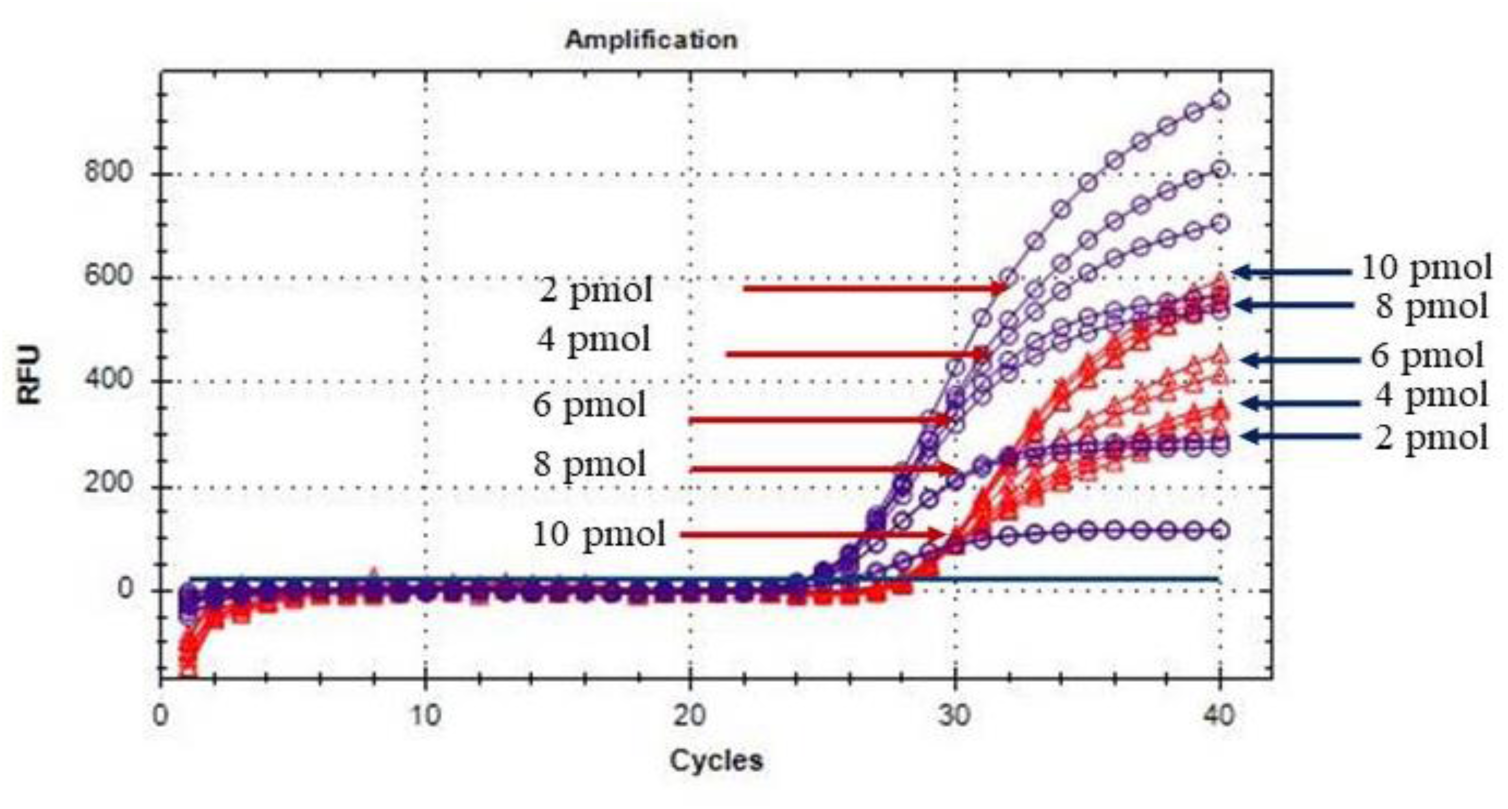
Titration of Primers and probes for *COWP* (FAM) indicating by red colour and *18ssu rRNA* (HEX) indicating by blue colour in duplex PCR method

**FIG 5a.**
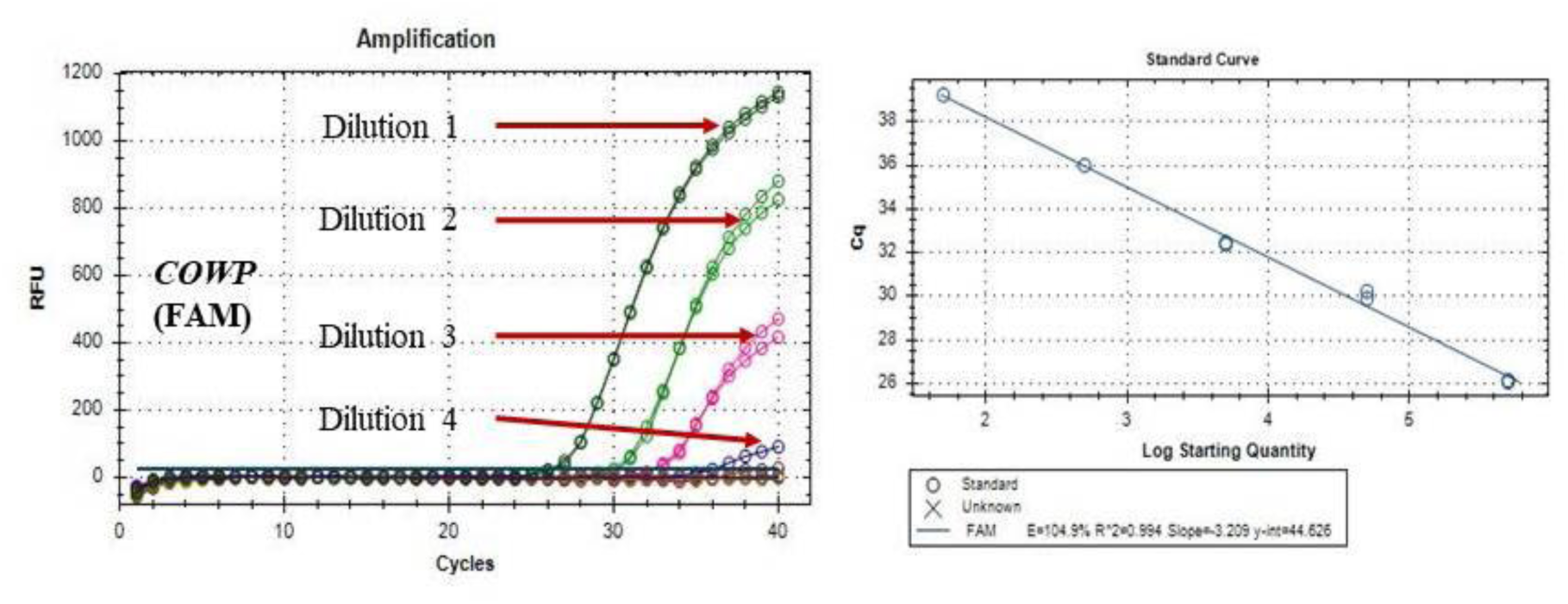
Amplification plot of serially diluted *Cryptosporidium* amplicon of *COWP-FAM* gene, descending concentrations (by a magnitude of Log_10_) ranging from 5.37×10^11^ (Dilution 1) to 7.08×10^05^ (Dilution6) for determination of LOD by TaqMan® probe based real-time PCR assay. 5**b** The standard curve for a 10-fold serial dilution series (with linearity from Dilution 1 to dilution 4) plotted as the threshold cycle on the Y-axis, against the target concentration of cDNA per assay (X-axis) with E value = 104.9%, correlation coefficient (R2) = 0.994, slope = −3.209 and Y intercept = 44.626

**FIG 6a.**
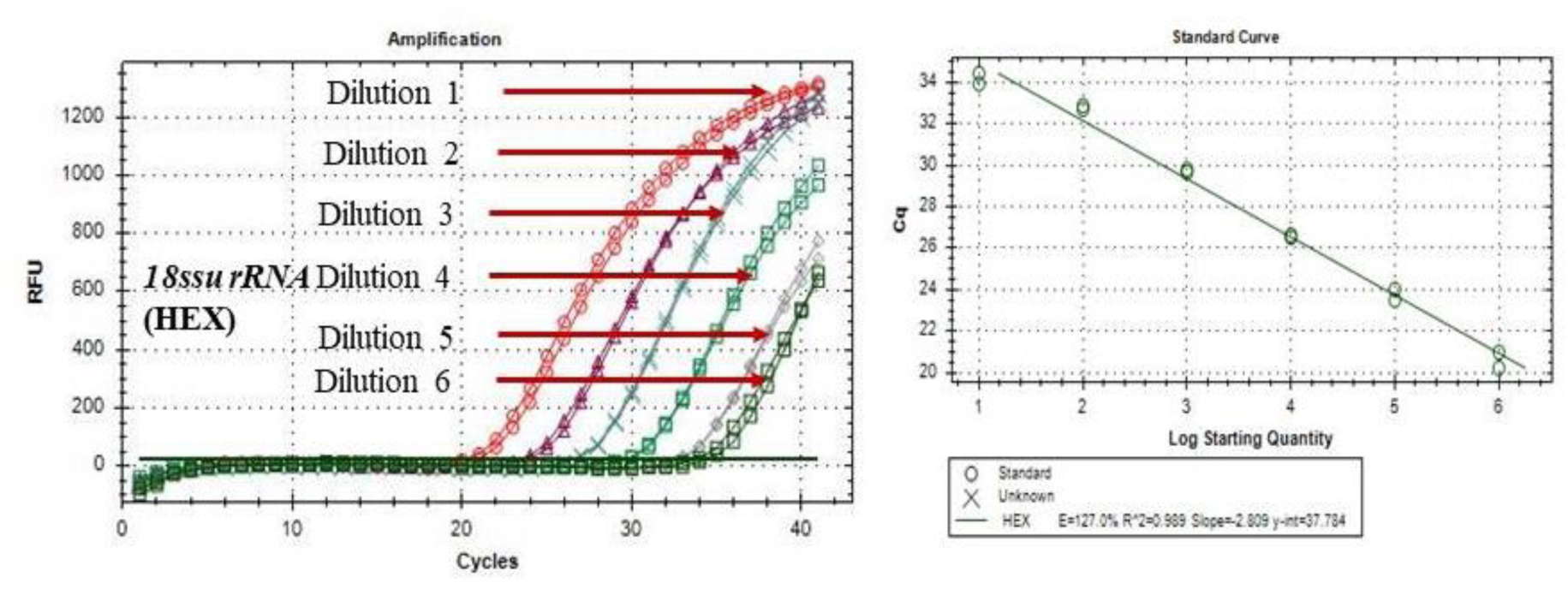
Amplification plot of serially diluted *Cryptosporidium* amplicon of *18ssu rRNA*-HEX gene, descending concentrations (by a magnitude of Log_10_) ranging from 1.22×10^12^ (dilution 1) to 1.22×10^07^ (dilution 6) for determination of LOD by TaqMan® probe based real-time PCR assay. **6b** The standard curve (with linearity from Dilution 1 to dilution 6) for a 10-fold serial dilution series plotted as the threshold cycle on the Y-axis, against the target concentration of cDNA per assay (X-axis) with E value = 127.0%, correlation coefficient (R2) = 0.989, slope = −2.809 and Y intercept = 37.784

**FIG 7a.**
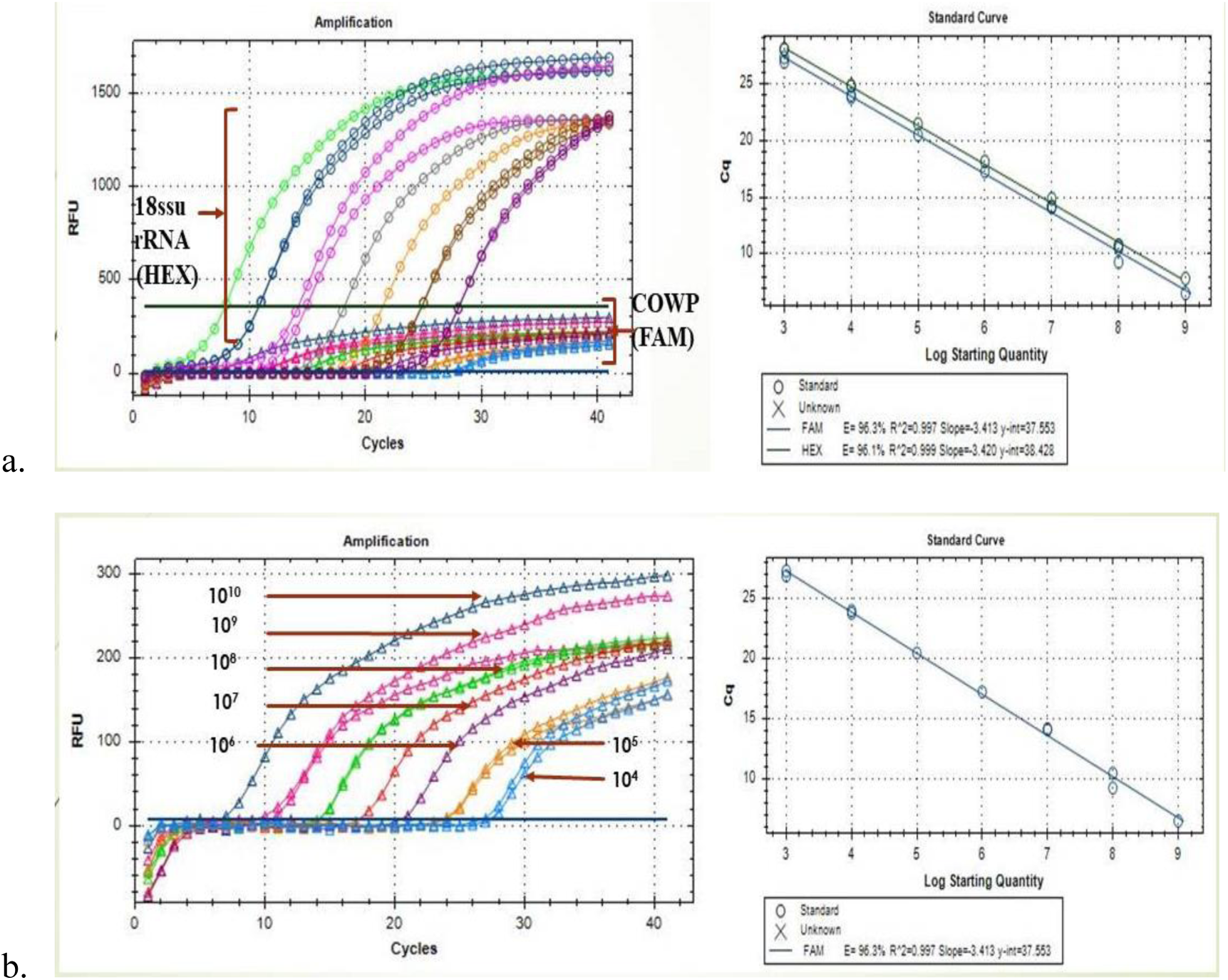
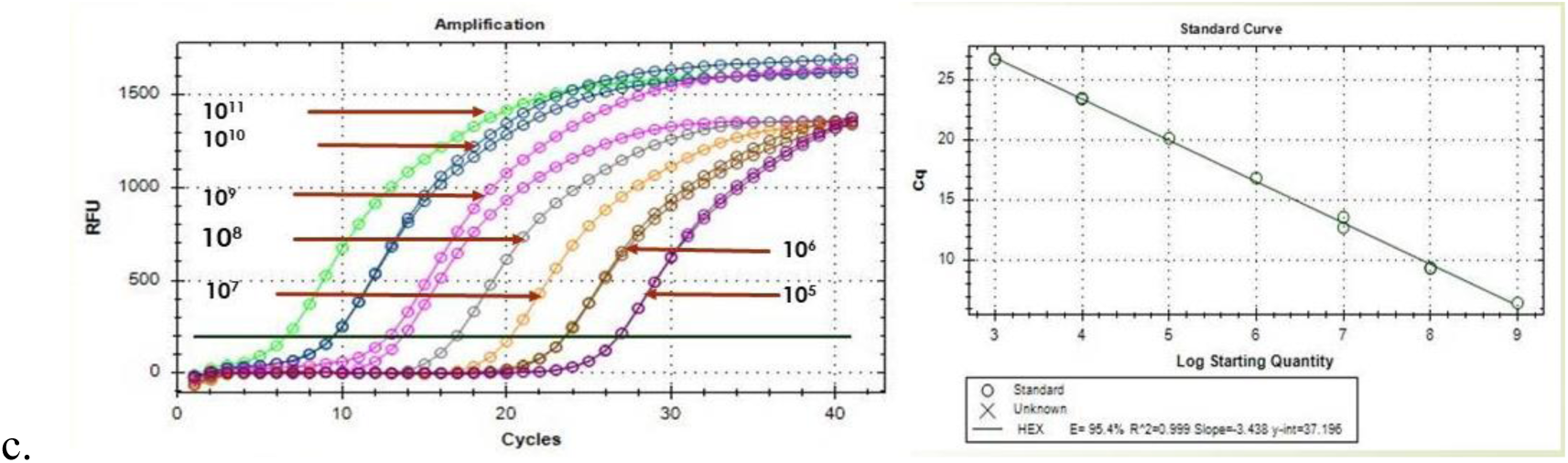
Duplex reverse transcription TaqMan^®^ Probe Real time PCR for *COWP* (FAM) and *18ssu rRNA* gene of *Cryptosporidium* from positive cDNA with standard curve, **7b** Extracted image of *COWP* gene only from the duplex real time PCR data and **7c** Extracted image of *18ssu rRNA* gene only from the duplex real time PCR data.

**FIG 8.**
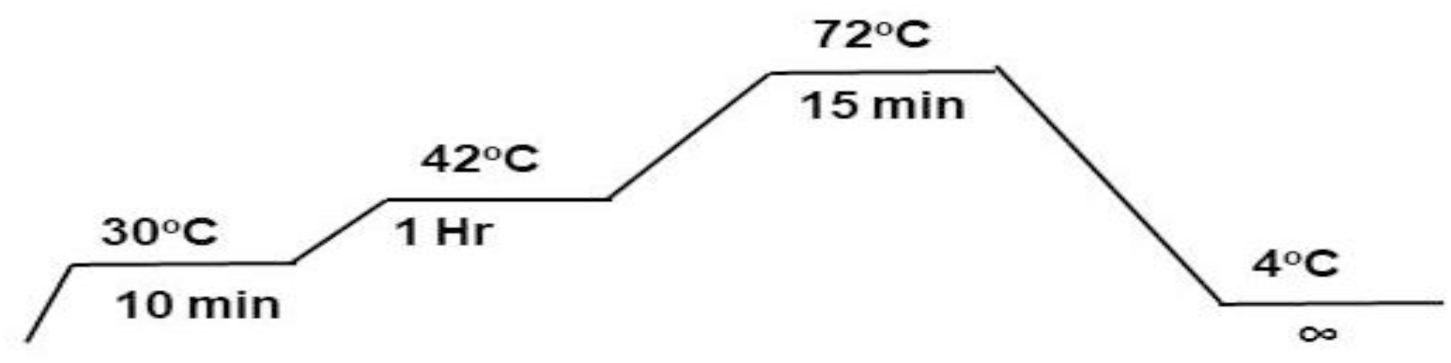
Cyclic conditions for cDNA synthesis in thermal cycler (Techne, TC 4000)

**FIG 9.**
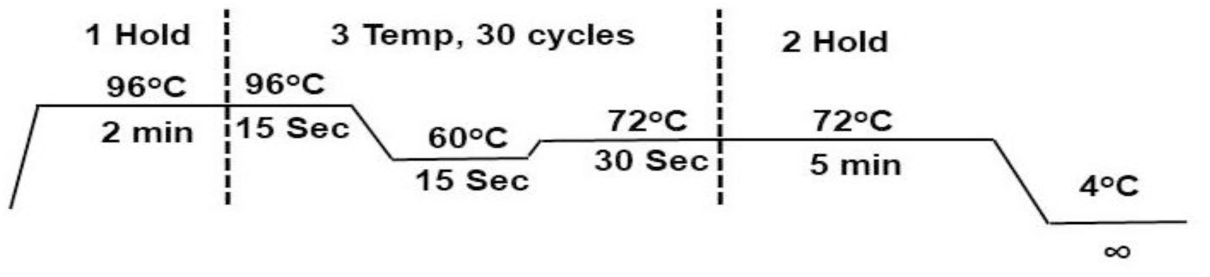
Cyclic conditions for gradient PCR in thermal cycler (Techne, TC 4000)

**FIG 10.**
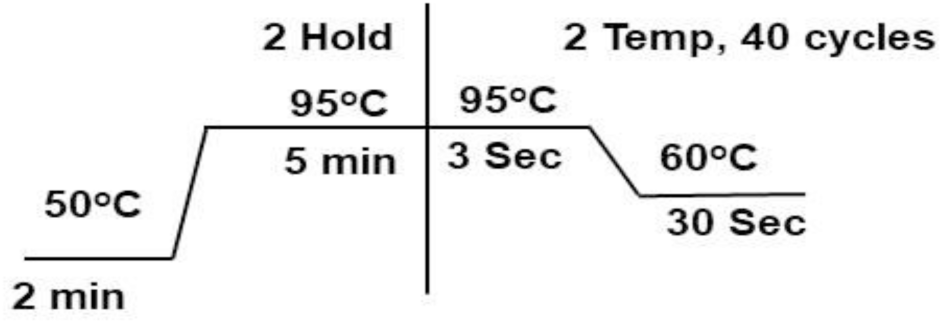
Cyclic conditions for DNA duplex TaqMan^®^ probe real time PCR

**FIG 1.**
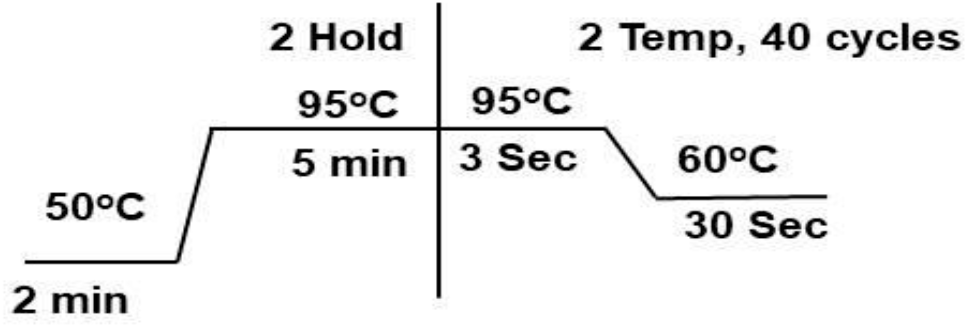
Cyclic conditions for primers and probe optimization protocol in TaqMan^®^ based real-time PCR (qPCR)

### Limit of Detection (LODs) using *COWP* (FAM) and *18ssu rRNA* (HEX), genes as simplex and duplex modes by reverse transcription TaqMan® probe real time PCR assay for *Cryptosporidium*

Limit of detection of target genes using simplex mode Based on the copy number calculation of respective purified amplicons of genes viz. *COWP* and *18ssu rRNA*, we performed limit of detection by serial dilutions assay in both simplex and duplex modes using the currently developed TaqMan probe real time PCR assay. Initially standard curves were generated for LOD by simplex assay.

The sensitivity of the assay was based on the copy numbers ranged from 5.37×10^11^ to 5.37×10^06^ for *COWP* (FAM) and 1.22×10^12^ to 1.22×10^07^ for *18ssu rRNA* (HEX) separately in simplex mode. The standard curve showed a linear curve over these dilutions to making the range of detection (ROD) for respective genes. The LOD for the targets *COWP*-FAM and *18ssu rRNA*-HEX were 5.37×10^06^ and 1.22×10^07^ copies respectively.

### Limit of detection of target genes using duplex mode

The sensitivity of duplex assay was based on the copy numbers ranged from 7.08×10^10^ to 7.08×10^05^ for *COWP* (FAM) and 5.95 ×10^11^ to 5.95 ×10^06^ for *18ssu rRNA* (HEX). The standard curve showed a linear curve over these dilutions to making the range of detection (ROD) for respective genes. The LOD for the targets *COWP*-FAM and *18ssu rRNA*-HEX were 7.08×10^05^ and 5.95 ×10^06^ copies respectively.

### Duplex reverse transcription TaqMan probe based Quantitative real time PCR (dRT-qPCR)

#### Relative quantification

The samples including unknown (clinically positive and cryptosporidia infected <1month olf goat kids) and control (Carrier status apparently healthy <1month goat kid positive for crytosporidial oocyst by microscopy) were assayed using Cryptosporidal dRT-qPCR along with NTC and NRT controls. The results and the fold change are given in the table below

## Discussion

Cryptosporidiosis is an important disease, which causes the diarrheal illness in neonatal animals associated with other diarrheal pathogens or as alone (32). However, it is difficult to diagnose cryptosporidiosis alone from other major etiologies of neonatal diarrhoea complex in neonatal animals including *E.coli*, Group A Rotavirus (GARV) and Bovine corona virus (BCoV). For diagnosing cryptosporidiosis in animals faecal oocyst identification by modified Ziehl-Neelsen staining acid fast staining is a preferred method. Although this method is enough to identify the acid fast oocysts based on colour, shape and size, but it cannot provide details of the active infective stages (33, 34). PCR is another method which is highly sensitive, specific and accurate (if standardized), but cannot fix the aforementioned problem of identification of active live oocysts from the passive or dead ones. Hence in the current study we have developed a reverse transcription based TaqMan probe duplex real time PCR for simultaneous detection of two target genes viz., COWP and 18ssu rRNA that would provide better detail on the active and passive stages of cryptosporidial infections in neonatal goats.

Microscopy based detection by modified Ziehl-Neelsen technique was conducted as a preliminary diagnostic screening for presence of cryptosporidial infection in suspected neonatal kids. As observed, this technique could identify 39 cases positive for Cryptosporidiosis out of 61 sampled. Microscopy is only a primary test, but gives an overall idea about the presence of cryptosporidiosis infection in the herd (21, 35–37). Therefore, many researchers used mZN technique as gold standard method for the screening of large number of population because it is much easier and non-invasive. Microscopy has certain limitations, like it may not differentiate between the active and passive infections and its sensitivity and specificity is lower as compared to ELISA and PCR (36, 38).

Specific diagnostic development with high accuracy and sensitivity is always on the priority to diagnose the each disease. Rapid diagnosis is an important way to overcome the impact of disease and appropriate treatment management. Diagnostic tests are inadequate, serological diagnosis, novel faecal diagnostic analysis and biomarker are require to develop the more accurate and sensitive identification of active *Cryptosporidium* infection (6).

In the present study, primer and probes were newly designed and checked for its working by using conventional PCR, followed by DNA based TaqMan probe duplex PCR. Both the tests identified the presence of Cryptosporidial DNA in the sample targeting the respective genes viz. *18ssu rRNA* and *COWP*.

Similarly, the Primers and probes were also titrated in simplex and duplex modes at various concentrations and optimised accordingly as given in the results. Eventually the optimal concentrations of primers and probes were used in the dRT-qPCR assay for unknown samples detection, standard curve using copy numbers and relative quantification based cryptosporidial transcriptional response. As per the copy number based standard curve, the RFU is higher for 18ssu rRNA because of the abundancy of the transcripts, while for the COWP gene the RFU is relatively lower and is present in clinically affected neonates with active cryptosporidial infection. Hence, there is a scope to decipher the active live cryptosporidal oocysts from the dead ones. Transcriptional activity is present only in live and metabolizing oocysts and their subsequent infective or reproductive stages (gametogony) and hence this assay has the ability to identify and at the same time quantify them based on gene targets used in the current study (34). The *18ssu rRNA* is a constitutively expressing gene which is present in all stages of life cycle of cryptosporidia, but transcripts for *COWP* gene is present only during specific stages and especially the active ones. There is a dearth of information as far as reverse transcriptional qPCR for cryptosporidiosis is concerned, because many researchers in the past used only DNA based detection method (29, 39) and most of them were used rather as a diagnostic for differentiation of *C.parvum* from *C.hominis* and other species (13, 30). Similarly sensitivity of detection is another parameter that decides the efficiency of the diagnostic assay, and previously researchers reported as low as 2 oocysts detected using TaqMan probe real time PCR assay and genus specific tests have better sensitivity compared to species specific assays (40). Recently DNA based commercial qPCR were tested for sensitivity and limit of detection, in which FTD stool parasites kit detected 1 and 10 oocyst for *C. parvum* and *C.hominis* respectively (41), while the in-house developed qPCR from the same research team showed up to 10 oocysts and 1000 oocysts for C. parvum and C.hominis respectively. Since most of these studies targeted the genomic DNA, we cannot compare them directly with our dRT-qPCR which is reverse transcriptional and targeting the RNA of the same pathogen. Also the mode of test was duplex in the current study unlike the previous researchers in which simplex mode was followed for LOD and sensitivity assays or multiplex assays involving other species like Giardia.

To further understand the dynamics of these two target genes, a relative quantification based dRT-qPCR assay was conducted. In this assay, the fold change was computed for *COWP* gene which is taken as a gene of interest (GOI) due to its differential expression during infections, while 18ssurRNA was used as a reference gene because of its constitutive expressive nature in cryptosporidial life cycle. Four clinically and microscopically cryptosporidia positive neonatal kids were compared with the control kids which are apparently healthy with no overt clinical signs but positive for cryptosporidia. This was done to actually assess the difference in expression levels of *COWP* in these two groups (clinical and carrier) of cryptosporidial positive animals. Also there is no logic in comparing the infected and non-infected animals, because here the genes are from the pathogen itself and not actually from the host.

Previously, PCR based test has been established for the identification of *Cryptosporidium* infection with higher sensitivity and specificity as compare to the conventional and other methods which used to identify the cause the infection in gastroenteritis (42–44). PCR based method gives the highly reliable and accurate results compare with faecal-smear microscopy because of cross-contamination from the environment in the direct faecal examination (45). Thus, PCR based test are helpful to identify the species or strain of any pathogens and differentiate the responsible pathogens for causing the diarrheal illness by their epidemiology and clinical manifestation (46). Therefore, this study used mRNA based TaqMan probe real time PCR assay to reliable and new diagnostic tool for detection the *Cryptosporidium* spp. at different stages of infection and differentiate the active and passive infection in host animal and humans.

The mRNA based TaqMan probe real time PCR assay is simple, rapid and self-regulating amplification system with low probability of cross-contamination as compared to other conventional PCR methods (47, 48). This study showed the successful standardization and development mRNA based TaqMan probe assay for targeting the *18ssu rRNA* and *cowp* genes.

## Material and Methods

### Sample collection

Diarrheic faecal samples were collected from Jamunapari and Barbari breed of goats from ICAR-CIRG Mathura. Faecal sample were collected as lavage from neonatal kids in 1X sterile PBS by flushing with the aid of sterile syringe (without any needle) through anal orifice (non-invasive method). The faecal lavage was transferred into a sterile 15ml falcon tube for further processing. The samples were centrifuged at 3500rpm for 30 minutes at room temperature. The Middle semisolid hazy layer is used to prepare faecal smear for Modified Ziehl-Neelsen staining (mZN staining) and nucleic acid extraction for molecular studies.

### Nucleic Acid extraction

#### RNA Extraction from Cryptosporidium

TRizol reagent (Cat# 9108) based protocol was used for RNA extraction of *Cryptosporidium.* For this 200µl of the processed faecal sample was reconstituted in 900µl of TRizol reagent in a 2ml of micro-centrifuge tube and vortexed at high speed for 30 seconds followed by incubation at 10 minutes for room temperature. The mixture is further spiked with 200µl chilled chloroform and briefly vortexed for 30 seconds followed by spin at 13000 rpm at 4°C for 15 minutes. The clear supernatant was collected in new micro-centrifuge tube (without touching the interface) and then equal volume chilled isopropanol is topped-up and allowed to precipitate for 30 minutes at 4°C. The sample mixture was centrifuged again at 13000 rpm for 15 minutes at 4°C. The supernatant was discarded without disturbing the pellet and reconstituted with 1ml chilled 70% ethanol to wash the pellet by centrifugation at 10000 rpm for 5 minutes at room temperature. Repeated this step for again and carefully discarded the supernatant and the residual solvent was aspirated with the aid of 10µl micropipette to obtain moist-free pellet. The Pellet is further air dried for 3-5 minutes at room temperature and reconstituted in 30μl of diethyl pyrocarbonate (DEPC) treated water. The freshly extracted RNA was quantified by mixing with Quantifluor RNA dye (Cat# E3310) using Quantus Fluorometer® (Promega Technologies, Madison, USA) as per the manufacturer’s protocol.

#### cDNA Synthesis

The quantified RNA was converted into copy DNA (cDNA) using Primescript™ 1^st^ strand cDNA synthesis kit (TaKaRa, Japan). 1^st^ Reaction Mixture was prepared by adding 8.0 µl of RNA (1.0μg/concentration adjusted to 125.0ng/µl), 1.0µl of Random Primer and 1.0µl of 10mM dNTP Mix and incubated at 65°C for 5 min and snap chilled in ice. 2^nd^ Reaction Mixture was prepared by adding 4.0µl 5x Primescript Buffer, 0.5µl RNAse inhibitor, 1.0µl PrimescriptRTase, 4.5µl RNAase free water in new micro-centrifuge tubes. The 1^st^ mixture the 2^nd^ mixture were mixed in a new 0.2ml PCR tube incubated in thermal cycler for reverse transcription at following conditions as per the thermal graph illustrated below.

### DNA Extraction from Cryptosporidium

#### DNA extraction

DNA extraction was also done for all the samples using the Wizard® Genomic DNA Purification Kit according to the manufacturer’s protocol (Cat#A1120, Promega USA).The eluted DNA was quantified by Quantus^TM^ Fluorometer® (Cat# E6150 Promega Technologies, Madison, USA) as per the manufacturer’s protocol and stored at -20 °C for further use.

#### Primer and probe designing

Real time PCR primers and probes (Table 1) were designed using the BioEdit software program version 7.0.5.3 (Hall, 2011), by aligning FASTA sequences of various available strains of Cryptosporidium sequences from GenBank (National Center for Biotechnology Information [NCBI]; http://www.ncbi.nlm.nih.gov/GenBank/). The primer sequences picked were further analysed using Oligo Analyzer software for binding and thermodynamic properties. Primer and probe sequences were then checked for cross-reactions with non-targeted sequences on the GenBank database using the Basic Local Alignment Search Tool (BLAST) (http://www.ncbi.nlm.nih.gov/blast/Blast.cgi) if any to rule out non-specific binding. Similarly probes were designed by the alignment of *COWP* gene and *18ssu rRNA* gene sequences according to the recommendations and general guidelines (Proudnikov et al., 2003). Reporter dyes FAM (Fluorescein amidites) and HEX (Hexachloro-fluorescein) were conjugated at 5’end of probe for *COWP* and *18ssu rRNA* genes respectively while BHQ -1 (Black Hole Quencher) used as quencher dye at 3’ end for both the probes. Primers and TaqMan® probes were synthesised commercially (Integrated DNA Technologies, Inc. 1710 Commercial Park Coralville, Iowa 52241 USA).

**Table 1.** Mean Cq and RFU values for COWP and 18ssu rRNA for dRT-qPRC

| S.No | Sample | *COWP* (Cq<br>Mean colored<br>blue) | Mean<br>RFU<br>values for<br>*COWP* | *18ssu rRNA*<br>Cq Mean<br>colored<br>green) | RFU values<br>for 18ssu<br>*rRNA* |
| --- | --- | --- | --- | --- | --- |
| 1. | Positive control | 24.84 | 393.56 | 21.58 | 696.75 |
| 2. | Sample 1 | 23.50 | 248.24 | 28.43 | 713.35 |
| 3. | Sample 2 | 28.31 | 239.46 | 26.42 | 626.51 |
| 4. | Sample 3 | 13.64 | 113.77 | 33.69 | 515.92 |
| 5. | NTC | 33.32 | 1.90 | 35.88 | 149.48 |

**Table 2.** primer probe optimization for duplex mRNA based TaqMan probe real time PCR

| S.N. | Genes | Final concentration used |  |
| --- | --- | --- | --- |
| 1. | *COWP* (FAM) | Forward Primer | 8pmol |
|  |  | Reverse Primer | 8pmol |
|  |  | Probe | 10pmol |
| 2. | *18ssu rRNA* (HEX) | Forward Primer | 4pmol |
|  |  | Reverse Primer | 4pmol |
|  |  | Probe | 4pmol |

**Table 3.** Concentrations used for serial dilutions in simplex LOD TaqMan probe assay

| S.No. | Gene | Stock<br>concentration | Starting<br>concentration | Ending<br>concentration | No. of<br>dilutions |
| --- | --- | --- | --- | --- | --- |
| 1. | *COWP* | 91ng/μl | $5.37 \times 10^{11}$ | $5.37 \times 10^{06}$ | 06 |
| 2. | *18ssu rRNA* | 144ng/μl | $1.22 \times 10^{12}$ | $1.22 \times 10^{07}$ | 06 |

**Table 4.** Concentrations used for serial dilutions in Duplex LOD TaqMan probe assay

| S.No. | Gene | Stock | Starting | Ending | No. of | Limit of detection |
| --- | --- | --- | --- | --- | --- | --- |

|  |  | concentration | concentration<br>(copy numbers) | concentration<br>(copy numbers) | dilutions | (copy numbers) |  |
| --- | --- | --- | --- | --- | --- | --- | --- |
|  |  |  |  |  |  | Upper limit | Lower limit |
| 1. | *COWP* | 12ng/μl | $7.08 \times 10^{10}$ | $7.08 \times 10^{01}$ | 10 | $7.08 \times 10^{10}$ | $7.08 \times 10^{04}$ |
| 2. | *18ssu*<br>*rRNA* | 70ng/μl | 5.95<br>$\times 10^{11}$ | $5.95 \times 10^{025}$ | 10 | 5.95<br>$\times 10^{11}$ | 5.95<br>$\times 10^{05}$ |

**Table 5.**
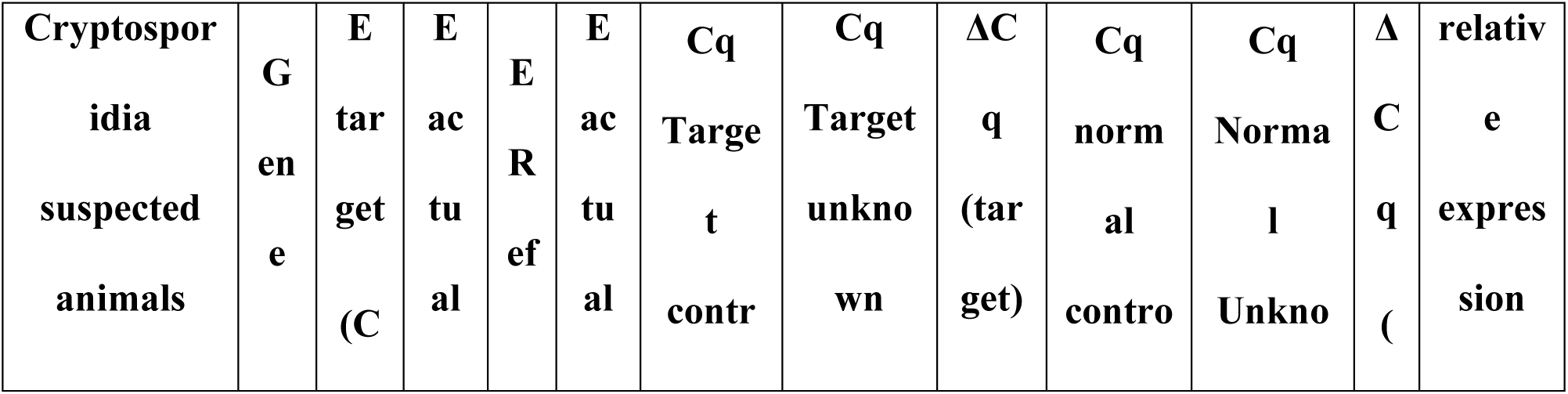
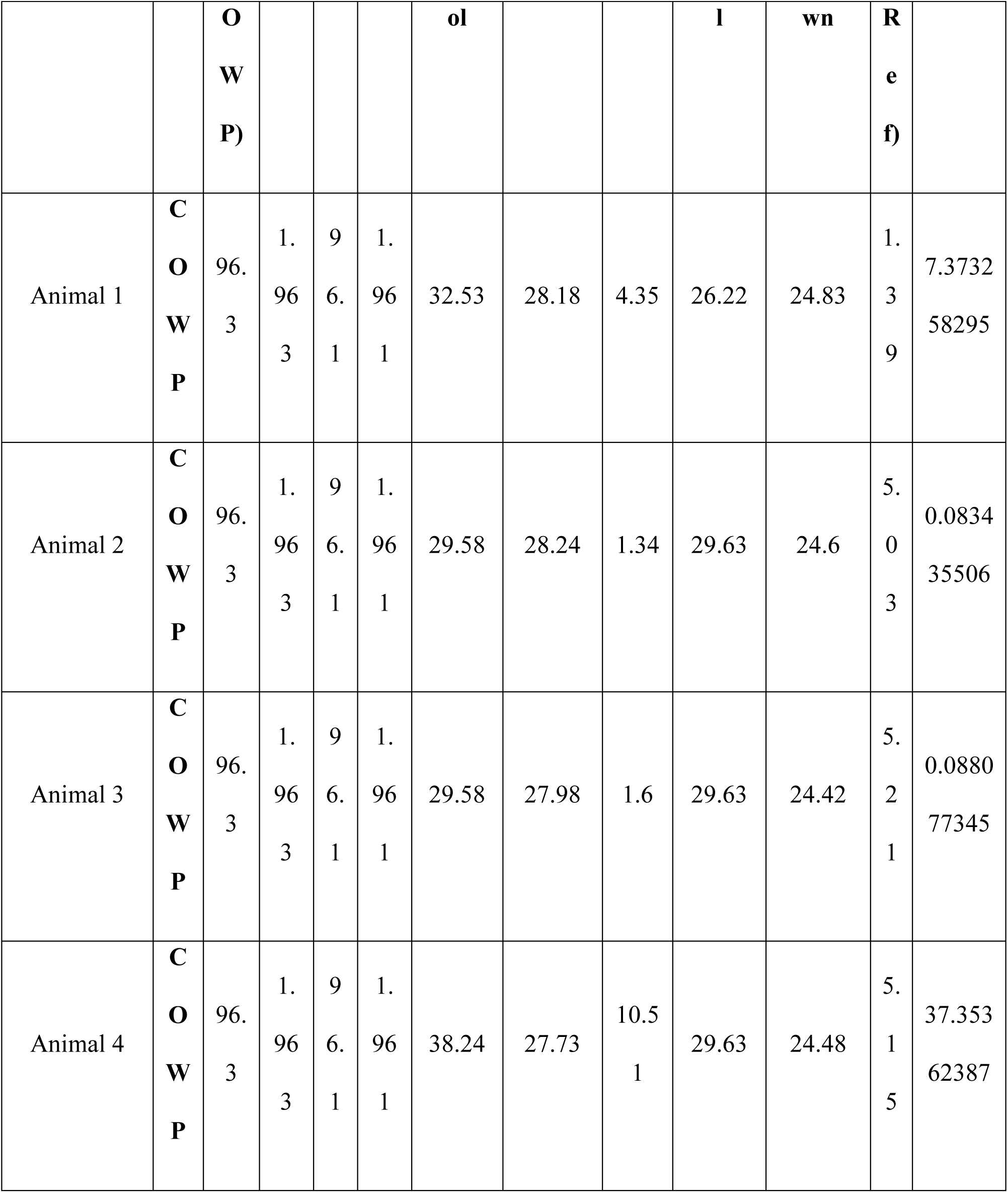
Relative quantification of *COWP* and *18ssu rRNA* by dRT-qPCR

**Table 6.**
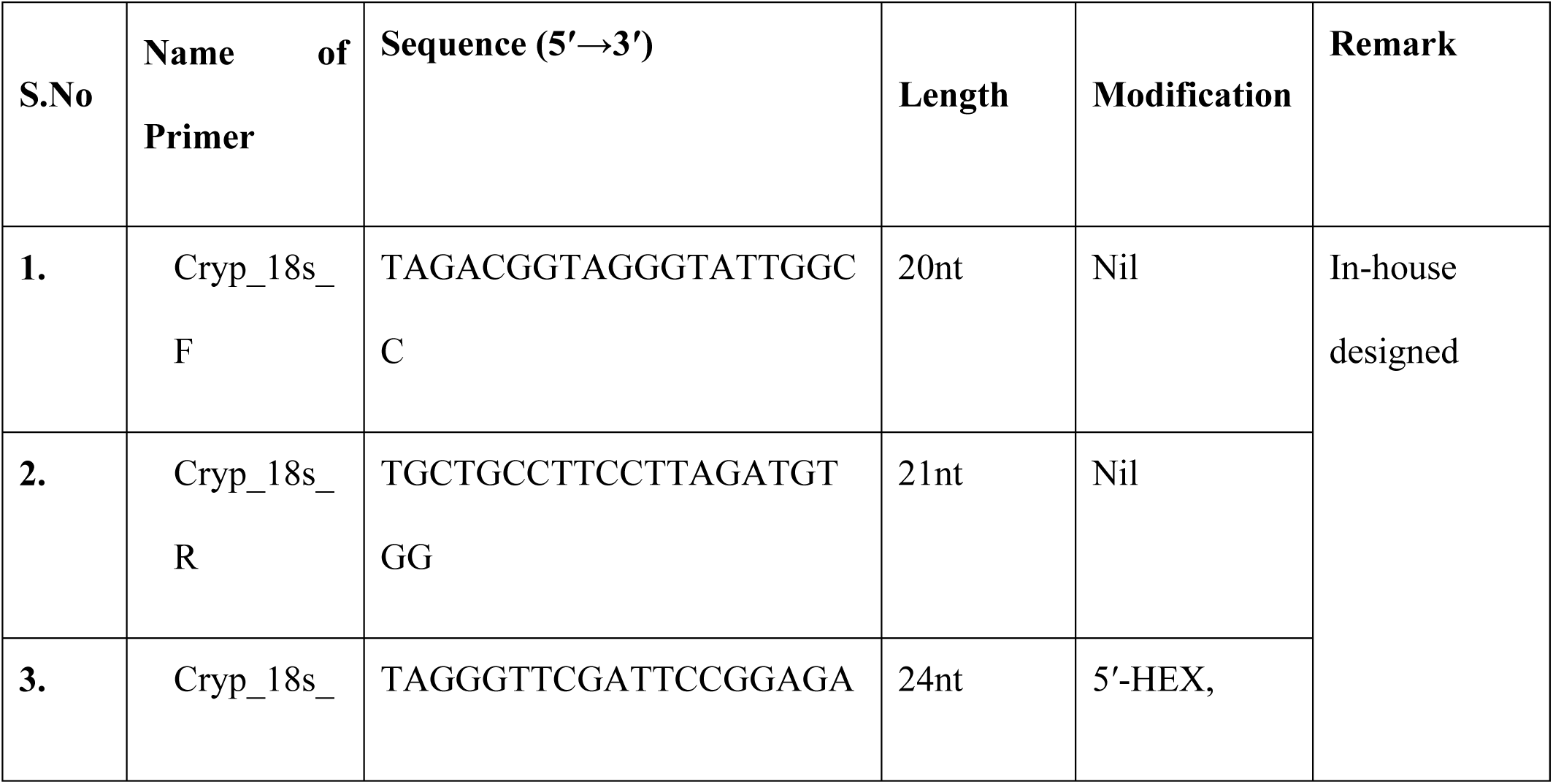
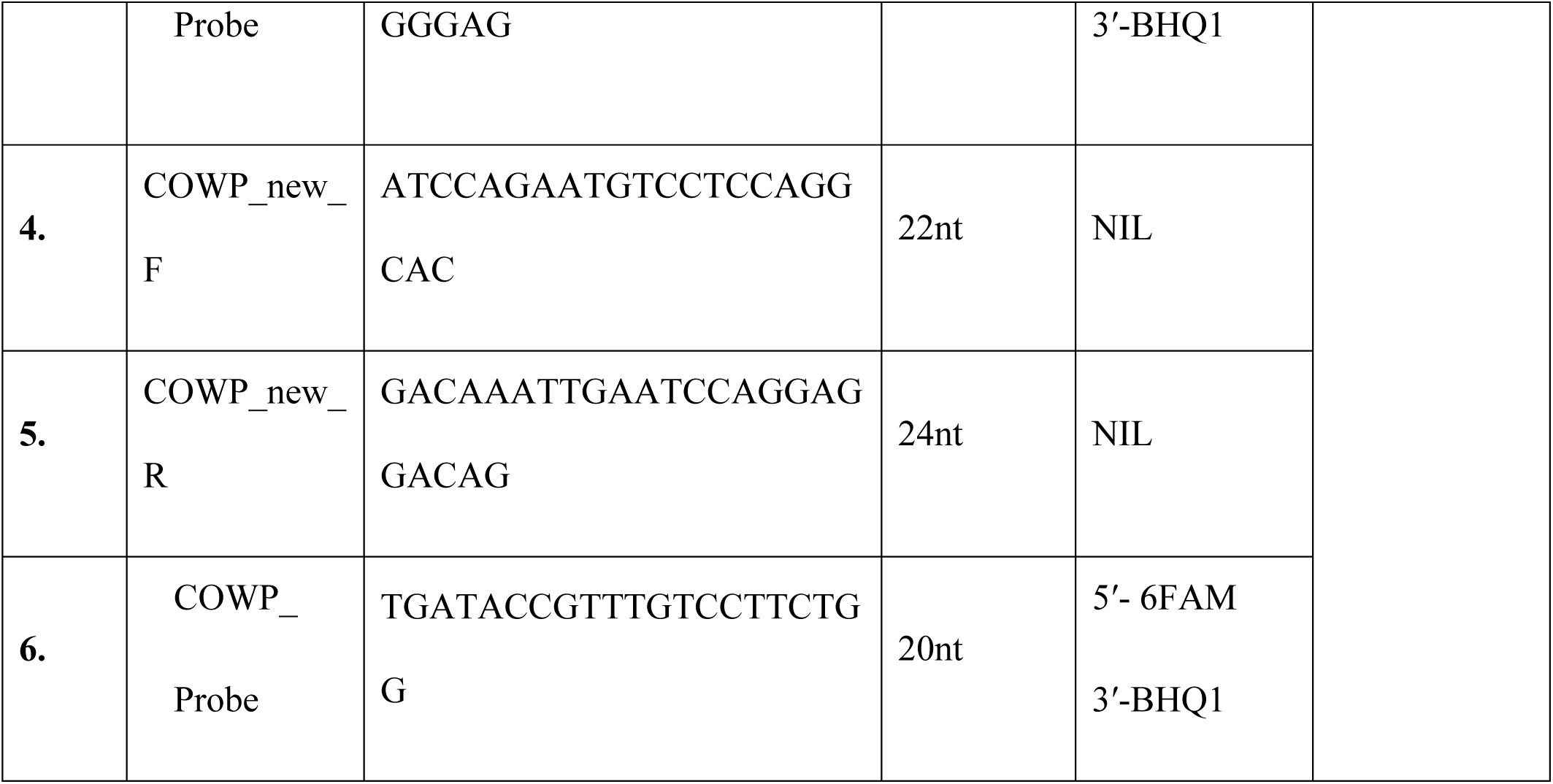
List of Primers and Probe used in TaqMan^®^ based real-time PCR

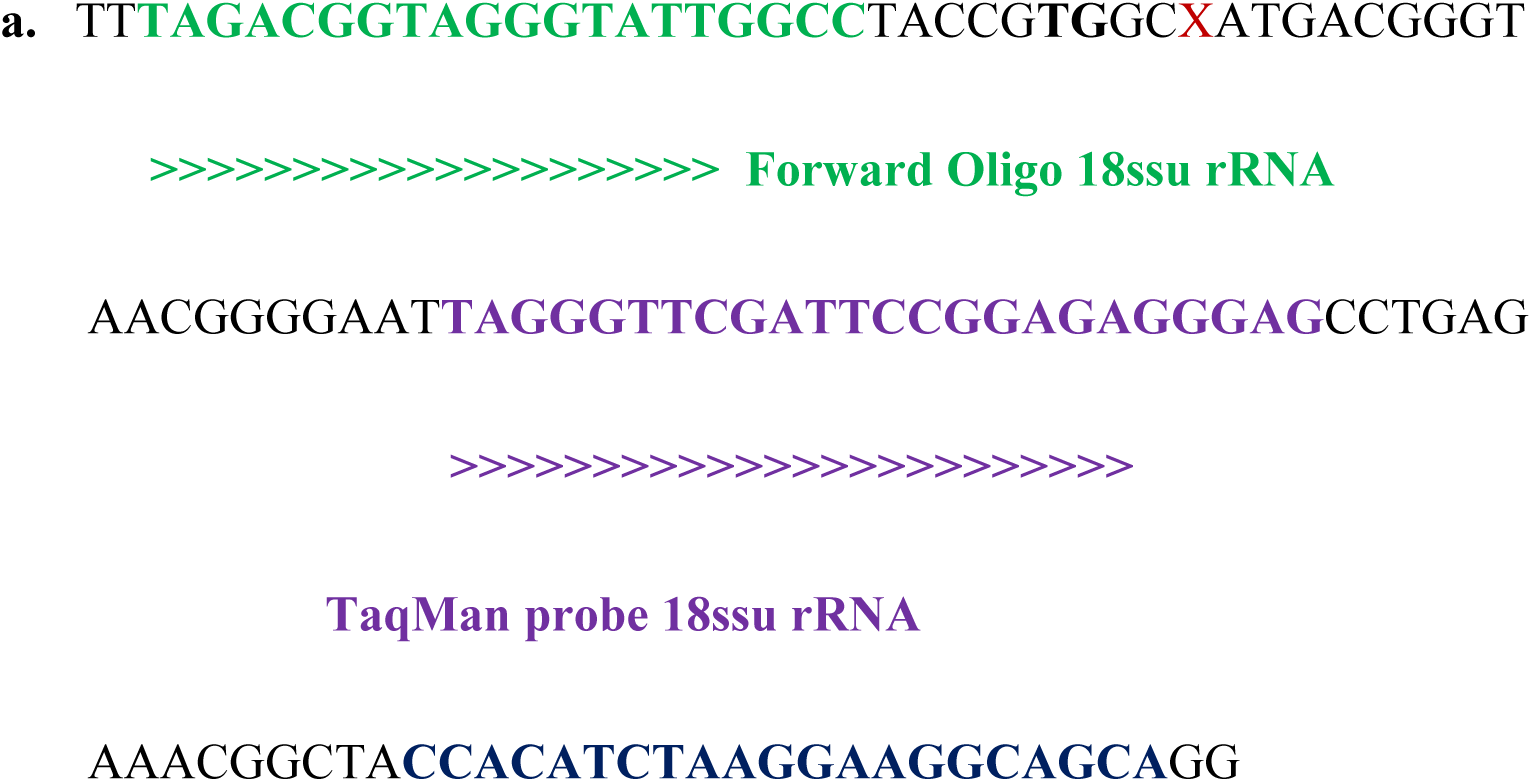
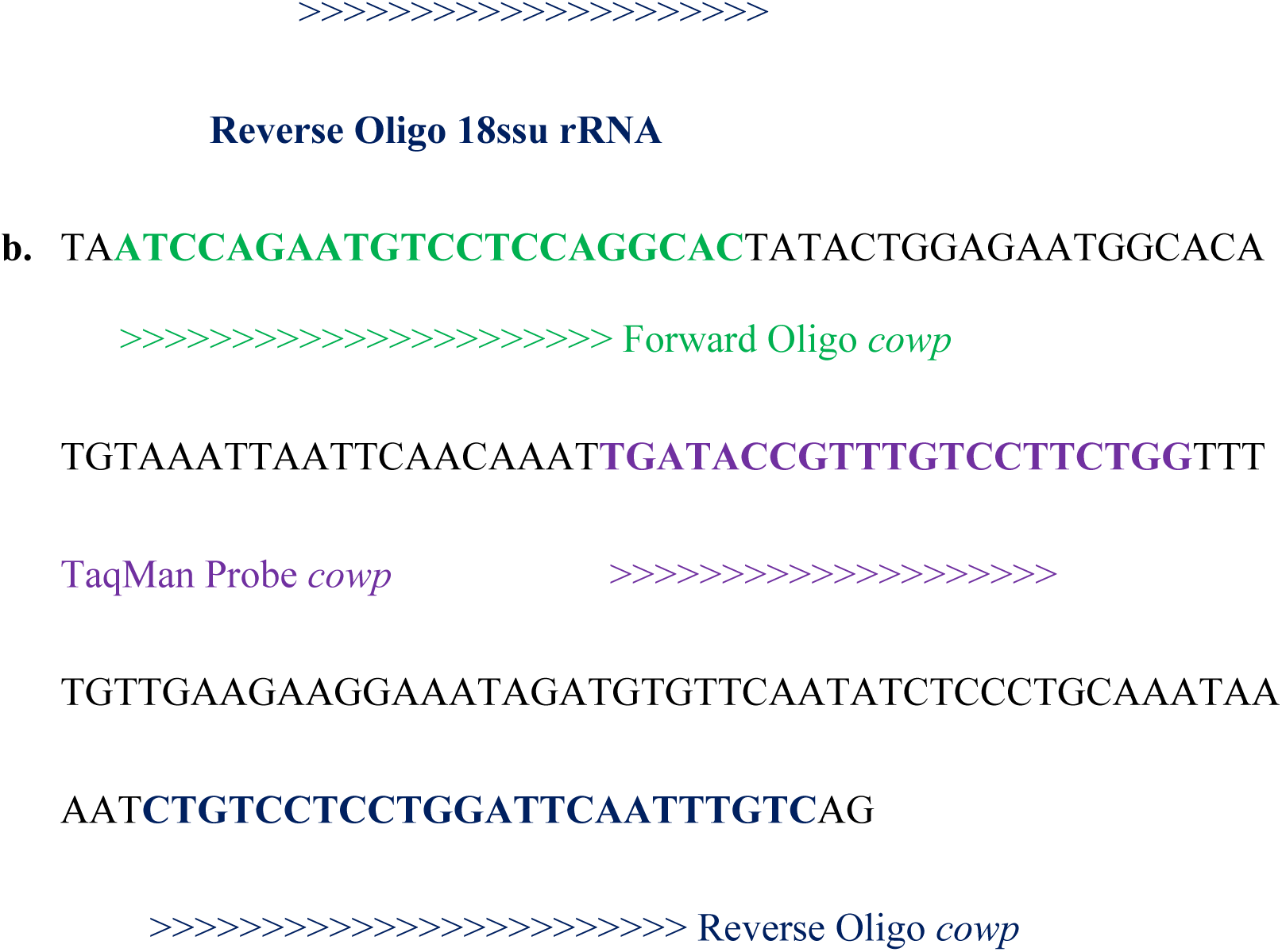

#### Conventional PCR

To check the workability each of the newly designed *18ssu rRNA* and *COWP* primers, conventional PCR was performed by following standard PCR reaction conditions. The conventional PCR was performed separately for the genes in a 25µl volume with 12.5µl 2x Emerald GT amp master mix (TaKaRa, Cat# RR310A) along with 10 picomole of each of the forward and reverse primers for the respective genes, 1µl of DNA template and made up the total volume with PCR grade water. All the components were mixed gently and amplification was carried out in thermal cycler (Techne, TC 4000) as per the thermal graph below.

Further the respective amplicons for the two genes were subjected to gel electrophoresis in 2.0 % agarose TAE gel stained with Ethidium bromide along with 100bp DNA ladder. Electrophoresis was carried out at a constant voltage of 100 V for 45 minutes. The DNA products were visualized under the UV trans-illuminator and photographed using gel documentation system.

### DNA duplex TaqMan**^®^** probe real time PCR

After confirmation by conventional PCR, the primers and newly designed respective probes (*18ssu rRNA* and *COWP*) were analysed for its proper binding using DNA-based duplex TaqMan® probe real time PCR. The real time PCR master mix was assembled in duplicates for each sample in a final reaction volume of 25µl in 8 strip PCR tubes with optically compatible caps assayed in the real-time thermal cycler (CFX96 Real time PCR system^®^, Bio-Radwith the reagent preparation (Table 7) and thermal conditions as mentioned below.

**Table 7.** Concentration and volume of reagents used in DNA duplex TaqMan^®^ probe real time PCR

| S.N<br>o | Reagent |  | Volume | Conc. used |
| --- | --- | --- | --- | --- |
| 1. | Premix Ex Taq Probe qPCR Master Mix |  | 12.5 µl | 2x |
| 2. | *COWP* (FAM) | Forward primer | 0.80 µl | (8 pmol/µl) |
|  |  | Reverse primer | 0.80 µl | (8 pmol/µl) |
|  |  | Probe | 1.0 µl | (10 pmol/µl) |
| 3. | *18ssu rRNA*<br>(HEX) | Forward primer | 0.4 µl | (4 pmol/µl) |
|  |  | Reverse primer | 0.4 µl | (4 pmol/µl) |
|  |  | Probe | 0.4 μl | (4 pmol/μl) |
| 5. | Template DNA |  | 3.0 μl | (~5ng) |
| 6. | Nucleus Free Water |  | 5.7 μl | - |
| Total |  |  | 25 μl |  |

### Standardization of Simplex Probe based reverse transcription real-time PCR

Real Time PCR protocol was separately standardized for *COWP* and *18ssu rRNA* probe and primers using positive control cDNA. Probe qPCR master mix (TaKaRa, Japan) was used for the quantitative analysis of *COWP* and *18ssu rRNA* genes. Primers and probes were titrated serially starting from 2pmol/reaction, 4pmol/reaction, 6pmol/reaction, 8pmol/reaction to 10pmol/reaction for *COWP* and *18ssu rRNA* genes individually (Table 8). All the reactions were performed in duplicates in Premix Ex Taq Probe qPCR Master Mix (2X) (TaKaRa, Japan Cat#RR390) in a final volume of 25µl per tube in a CFX96 Real time PCR system^®^ (Bio-Rad) with the thermal conditions as described below. The reaction also included positive control (PC) and No template Control (NTC) for interpretation of the results.

**Table 8.** Optimization and standardization of Probe and primer dilutions with checkerboard titration method for Simplex real time PCR of *COWP* and 18ssu *rRNA* genes

| Targeted gene | Dilution |  |  |  |  |  |
| --- | --- | --- | --- | --- | --- | --- |
| *cowp*<br>Probe<br>dilution | Forward and Reverse<br>Primer<br>(constant) | 10 pmol each as constant |  |  |  |  |
|  | Probe (titration) | 2pmol | 4pmol | 6pmol | 8pmol | 10pmol |
| *cowp*<br>Primer<br>dilution | Forward and Reverse<br>Primer<br>(titration) | 2pmol | 4pmol | 6pmol | 8pmol | 10pmol |
|  | Probe<br>(constant) | 10 pmol constant |  |  |  |  |
| *18ssu rRNA*<br>Probe<br>dilution | Forward and Reverse<br>Primer<br>(constant) | 10 pmol each as constant |  |  |  |  |
|  | Probe (titration) | 2pmol | 4pmol | 6pmol | 8pmol | 10pmol |
| *18ssu rRNA*<br>Primer<br>dilution | Forward and Reverse<br>Primer<br>(titration) | 2pmol | 4pmol | 6pmol | 8pmol | 10pmol |
|  | Probe<br>(constant) | 10 pmol constant |  |  |  |  |

**Table 9.** Concentration and volume of reagents used in Probe based real-time PCR

| S.No | Reagent | Volume | Conc. Used |
| --- | --- | --- | --- |
| 1 | Premix Ex Taq Probe qPCR Master Mix | 12.5 µl | 2x |
| 2 | Forward primer | 1.25 µl | (10 pmol/µl) |
| 3 | Reverse primer | 1.25µl | (10 pmol/µl) |
| 4 | Probe | 1.25 µl | (10 pmol/µl) |
| 5 | Template | 1.0 µl | (~5ng) |
| 6 | Nucleus Free Water | 7.75 µl | - |
|  | **Total reaction volume** | **25µl** |  |

**Table 10.**
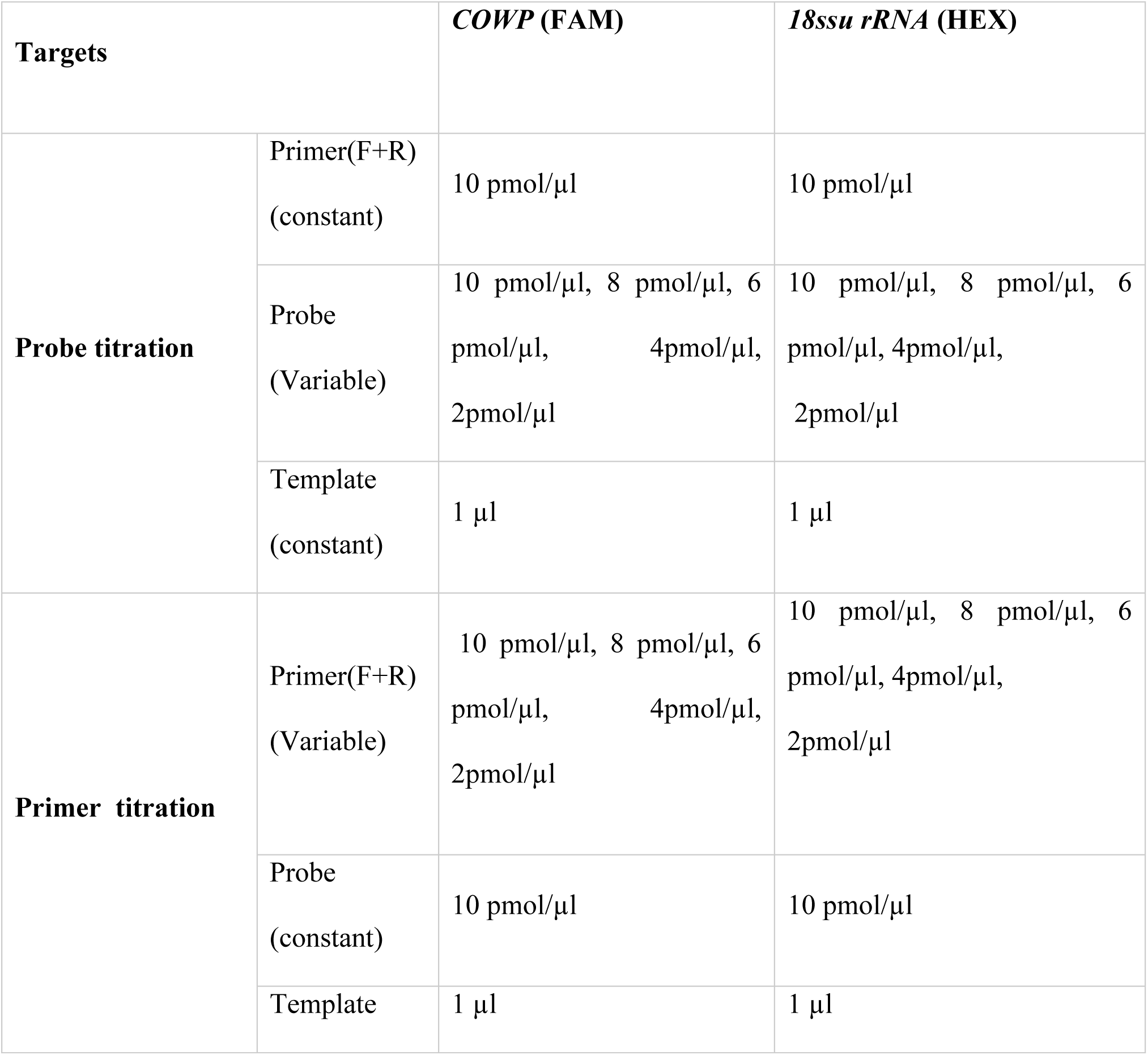
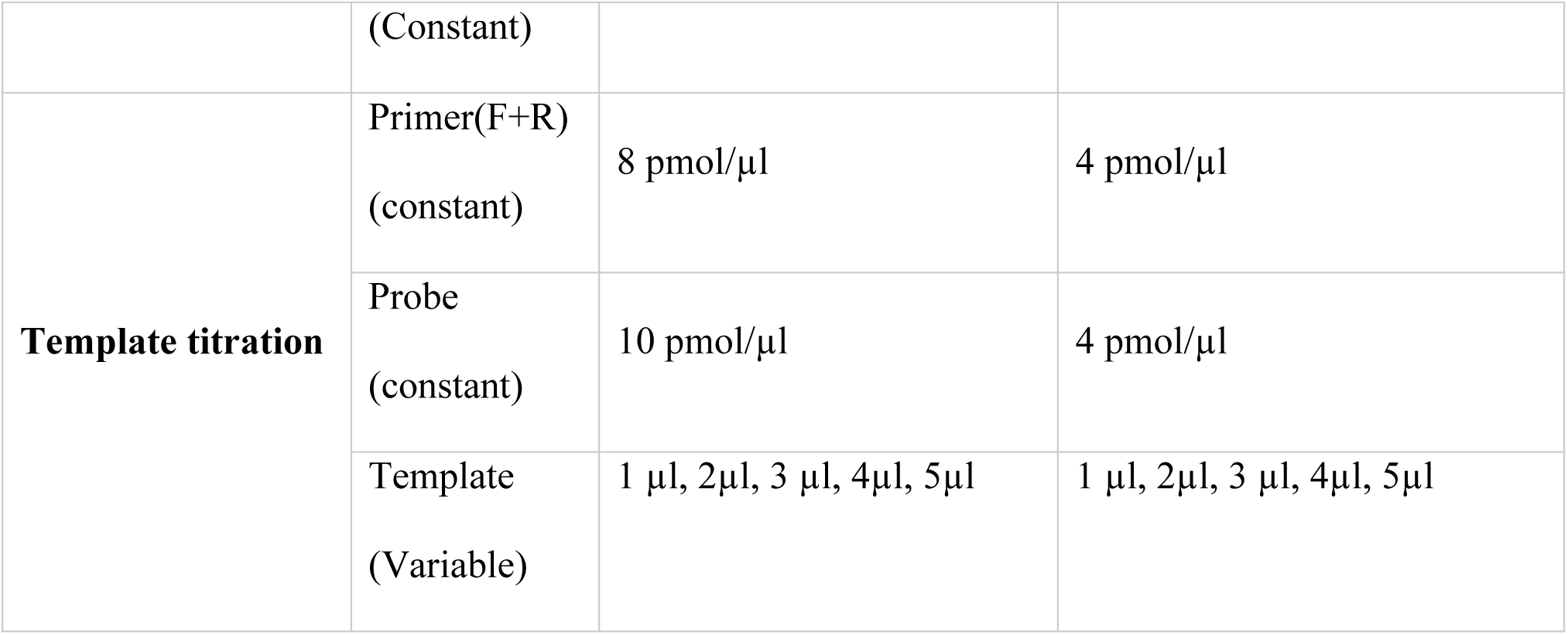
Optimization of primers, probes and template dilutions with respective constants and variables concentration for multiplex real time PCR of all three parameters and both the genes.

**Table 11.**
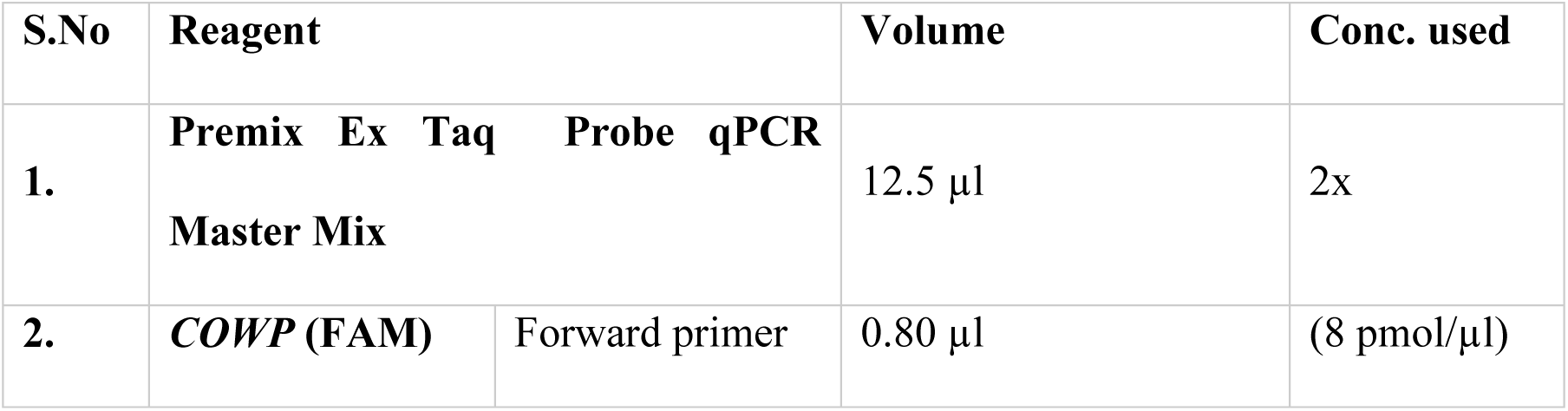
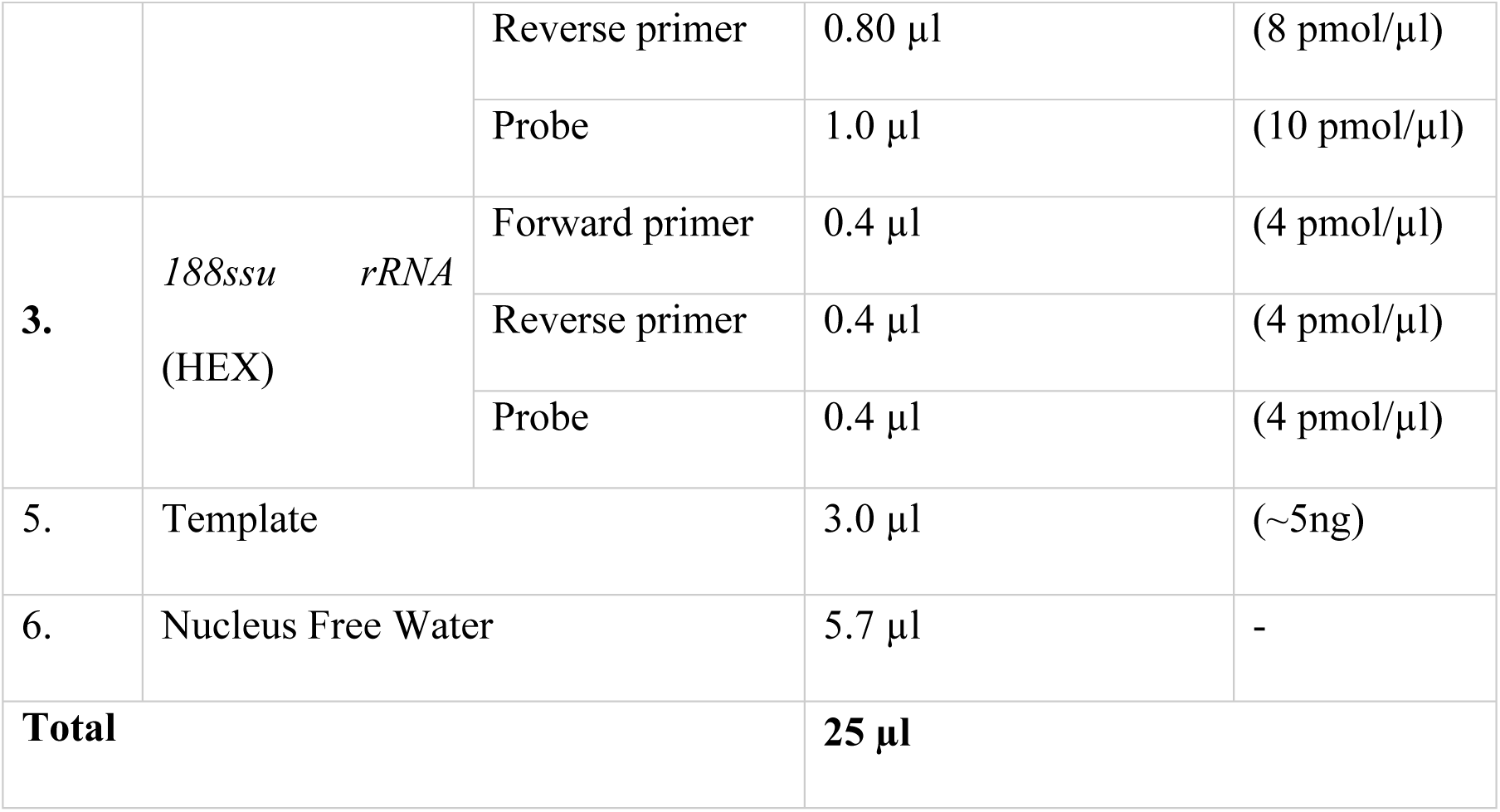
Final concentrations of primer, probe and templates in multiplex TaqMan^®^ probe based Real Time PCR reaction mix

### Duplex probe Real time PCR

#### Standardization and optimization of primer, probe & template for Duplex reverse transcription TaqMan^®^ Probe Real time PCR for *Cryptosporidia*

Finally, after conducting a series of experiments on the primer and probe standardization, the same assay was subjected to duplex mode using RNA as target. As per the previous section on DNA based duplex PCR, a similar titration technique is followed but this time with positive cDNA prepared from oocysts of *Cryptosporidium*.

#### Duplex reverse transcription TaqMan^®^ Probe Real time PCR for *Cryptosporidia* after optimization

After standardization and optimization Duplex reverse transcription TaqMan**^®^** Probe Real time PCR for *Cryptosporidia* was performed with total reaction volume of 25µl with the same thermal cyclic conditions as above. Final reaction concentration of primer, probe and template for *COWP* (FAM) and *188ssu rRNA* (HEX) genes are given in table below.

#### Limit of detection assays for the Duplex Probe Real time PCR

To know the limit of detection (LOD) and other variables including regression coefficient, Y-intercept, efficiency etc, of the current assay, a copy number based standard dilution was conducted by absolute quantification mode. Positive DNA amplicons generated by conventional reverse transcription PCR for *COWP* and *18ssu rRNA* genes respectively were gel purified using Sigma GenElute gel extraction kit (Cat# NA1111) following manufacturer’s protocol. The gel purified DNA were further subjected to quantification by mixing with QuantiFluor ONE dsDNA System, (Cat# E4871 Promega) and measured in Quantus^TM^ Fluorometer® (Cat# E6150, Promega) as per the manufacturer’s protocol. Copy number was derived based on the amplicon length (bp) and concentration of the purified amplicon by using the copy number formula **available in the** online tool (https://cels.uri.edu/gsc/cndna.html) as given below 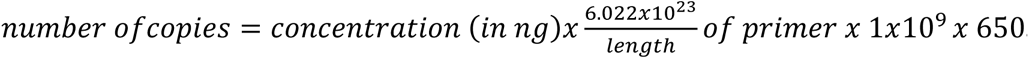. The real time PCR conditions and master mix assembled for the LOD were the same as mentioned in the previous section for duplex TaqMan probe real time PCR.

#### Duplex reverse transcription TaqMan probe based Quantitative real time pcr (dRT-qPCR) Relative quantification

Relative quantification was performed to quantify the transcriptional fold change of COWP (Target gene) vis-à-vis the 18ssurRNA gene (Reference gene). Since 18ssurRNA gene was constitutively expressed in cryptosporidia, it was regarded as a reference or housekeeping gene in the current study. For this relative quantification assay, 4 numbers of neonatal goat kids of less than one month of age were selected (based on screening from the 100 animals stock of same age group) for this study based on clinical signs (clinical status) like diarrhea, weakness and dehydration along with detection of positive oocysts in fecal mZN smear microscopy. Similarly, a neonatal goat kid of same age which is apparently healthy (no signs of diarrhea, dehydration, weakness, dullness etc.) but positive for cryptosporidial oocysts (carrier status) by fecal microscopy has been selected as control animal for the study based on continuous screening of healthy animals from a stock of 100 animals of same age group (<1month age). Further fecal samples from the kids with clinical status (Unknown) and carrier status (Control) were subjected to RNA extraction and cDNA synthesis as per the protocol described in the previous sections. Reaction mix was prepared in duplicates for each of the sample using the master mix, probes, primers, and nuclease free water as per the protocol described in previous sections (Table.No.6) in duplex mode. No template control (NTC) and non-reverse transcription controls (NRT) were also kept for interpretation of the results. The assay was performed as per the thermal conditions described in previous sections (Fig.4). The data analysis for gene expression studies was done using the CFX-96 manager (CFX Real-time PCR system, Biorad^®^, USA) using the ΔΔCq algorithm. The ΔΔCq method was done using the formula described by Livak and Schmittgen (2001) for determining the relative expression (fold change) of target genes of cryptosporida in total RNA isolated from fecal samples of neonatal goat kids used in the current study. In this method, the ΔCq (cycle threshold) values for each cryptosporidia suspected goat kids were figured out by subtracting 18ssurRNA Cq values from Cq values of gene of interest (in our study it is COWP). The average ΔCq for samples from control group (healthy cryptosporidia oocyst carriers with no clinical signs) are used as calibrator (ΔCq calibrator).

Further ΔΔCq values were computed by subtracting the average ΔCq of calibrator (healthy cryptosporidia oocyst carriers with no clinical signs) group from average ΔCq of unknown animals (clinically and microscopically positive for cryptosporidia).

ΔΔCq = ΔCq (Unknown) –ΔCq (control)

where, ΔCq (unknown) = Cq (target gene; COWP) – Cq(reference gene; 18ssurRNA)

Therefore,

ΔΔCq= [Cq(target, unknown)−Cq(ref, unknown)]−[Cq(target, control)−Cq(ref, control)]

Based on the above formula the fold change or relative expression for COWP gene was calculated by taking ΔΔCq as negative exponent of two, i.e., 2^-ΔΔCq^. The fold change thus derived represents the gene expression normalized to an endogenous reference or house-keeping gene (18ssurRNA) and relative to the control. A fold change of 1 in control group was used as a baseline for comparison with unknown animals (clinically and microscopically positive for cryptosporidia).

